# Papillary renal cell carcinomas rewire glutathione metabolism and are deficient in anabolic glucose synthesis

**DOI:** 10.1101/651265

**Authors:** Ayham Alahmad, Vanessa Paffrath, Rosanna Clima, Jonas Felix Busch, Anja Rabien, Ergin Kilic, Sonia Villegas, Bernd Timmermann, Marcella Attimonelli, Klaus Jung, David Meierhofer

**Author notes:** Communicating author – contact information: David Meierhofer – Max Planck Institute for Molecular Genetics, Mass Spectrometry Facility, Ihnestrasse 63-73, 14195 Berlin, Germany.

## Abstract

Papillary renal cell carcinoma (pRCC) is a malignant kidney cancer with a prevalence of 7-20% of all renal tumors. Proteome and metabolome profiles of 19 pRCC and patient-matched healthy kidney controls were used to elucidate the regulation of metabolic pathways and the underlying molecular mechanisms. Glutathione (GSH), a main reactive oxygen species (ROS) scavenger, was highly increased and can be regarded as a new hallmark in this malignancy. Isotope tracing of pRCC derived cell lines revealed an increased *de novo* synthesis rate of GSH, based on glutamine consumption. Furthermore, rewiring of the main pathways involved in ATP and glucose synthesis was observed at the protein level. In contrast, transcripts encoding for the respiratory chain were not regulated, which prompts for non-genetic profiling. The molecular characteristics of pRCC are increased GSH synthesis to cope with ROS stress, deficient anabolic glucose synthesis, and compromised oxidative phosphorylation, which could potentially be exploited in innovative anti-cancer strategies.

**SIGNIFICANCE STATEMENT:** We applied proteome- and metabolome profiling to elucidate molecular features in malign papillary renal cell carcinomas. By this characterization, a reprogramming of the main metabolic pathways, such as gluconeogenesis and fatty acid- and amino acid metabolism were identified. The proteins involved in the respiratory chain and the corresponding enzymatic activities were strongly reduced in pRCC, showing an anti-correlation compared with the transcriptome. Similar to renal oncocytomas, the ROS scavenger glutathione was identified as a hallmark in pRCC. Our results suggest that impaired metabolism and dysfunctional mitochondria determine the fate of pRCC. Furthermore, we propose that the specific regulation of the mitochondrial respiratory chain can differentiate highly similar malignant pRCCs from benign renal oncocytomas.

## INTRODUCTION

Papillary renal cell carcinoma (pRCC) is a heterogeneous disease, representing 7-20% of all renal cancers (1–3), subdivided into clinically and biologically distinct type I and type II entities (3). Type I pRCC tumors consist of basophilic cells with papillae and tubular structures and small nucleoli, whereas pRCC type II tumors exhibit large cells with abundant eosinophilic cells and prominent nucleoli (3). The cytogenetic differences are a trisomy 7 and 17 in pRCC type I and the loss of 1p and 3p in pRCC type II tumors (4, 5). Intra- and interchromosomal rearrangements are significantly increased in pRCC type II, leading frequently to a gene fusion involving the transcription factor *TFE3* (6), by which the promoter substitution appears to be the key molecular event, causing dysregulation of many signaling pathways already implicated in carcinogenesis (7).

Type I of this malignant tumor is characterized by frequent mutations in the *MET* oncogene, including alternative splice variants. Currently, the MET pathway is the most common target for developing new treatments for pRCC, such as the MET kinase inhibitor Savolitinib, which interrupts angiogenesis (8). Type II has more likely mutations such as *SETD2*, *NF2*, and the inactivation of *CDKN2A* by mutation, deletion, or CpG island hypermethylation (5, 9). Structural variants were observed in sporadic events, including duplications in *EGFR* and *HIF1A*, and deletions in *SDHB, DNMT3A*, and *STAG2* (10). These genes and several more, which can be found mutated in both tumor types, play a pivotal role in epigenetic regulation, signaling, and proliferation regulation, such as in PI3K/AKT/mTOR, NRF2-ARE, and the Hippo pathways. Furthermore, type II pRCC was subdivided into three subtypes, based on distinct molecular and phenotypic features. pRCC type II tumors are more likely to metastasize (4), and *FH* mutations and DNA hypermethylation were found to be correlate with inferior prognosis (5, 9). Hence, the hypermethylation group was termed “CpG island methylation phenotype” (CIMP), which additionally featured a metabolic shift known as the Warburg effect (5).

Many studies have been performed recently at the transcript level in pRCC to better understand its classification and subclassification and to elucidate pathway remodeling in these cancer (5, 11) studies. Based on all these findings, a new classification system was proposed. Compared with the current model where the organ of origin determines the tumor type, a system was proposed based solely on molecular features which could be considered more relevant for tumor classification (12).

Nephrectomy or partial nephrectomy and in the presence of metastases, and treatment with VEGF and mTOR inhibitors are currently considered the standard treatments. Furthermore, resection or irradiation of metastases can be a useful palliative treatment for patients with brain metastases or osseous metastases that are painful or increase the risk of fracture (13).

Besides transcript data, little is known in pRCC about its regulation at the protein- and metabolome level, the underlying molecular mechanisms, the alterations of metabolic pathways, and how well these “omics” data correlate with each other. One study compared metabolomic/lipidomic profiles of clear cell RCC (ccRCC), chromophobe RCC (chRCC), and pRCC, and determined that RCC subtypes clustered into two groups separating ccRCC and pRCC from chRCC, which mainly reflected the different cells of origin (14).

To fill in the aforesaid knowledge gaps, we undertook a multi-‘omics’ survey to compare seven type I, seven type II, and five metastatic type II pRCCs with patient-matched adjacent healthy kidney tissues. To confirm the main findings obtained by profiling, the dysregulated pathways were validated by either enzymatic measurements or isotope tracing experiments.

## RESULTS

### Proteome Profiling of pRCC

Malignant and non-malignant tissues from 19 nephrectomies representing papillary RCC of type I, II, and IIM (with metastases) were investigated; clinical parameters are shown in Table 1. Proteome profiling revealed a total of 8,554 protein groups, consisting of 1,330,129 identified peptides in all 19 pRCC samples and adjacent healthy kidney tissues, both at a false discovery rate (FDR) of 1% (Dataset S1). 3.785, 3.838, and 4.200 protein groups could be quantified by label-free quantification in type I, II, and IIM, respectively. Pearson correlation ranged from 0.597 to 0.951 for controls and 0.631 to 0.938 for all pRCC specimens for the least to the most similar individuals. Furthermore, proteome profiling revealed pRCC I, II, and IIM versus healthy adjacent kidney tissues as distinct groups in a principal component analysis (Figure 1A-C). Significantly regulated proteins were identified by a t-test and volcano plots are shown for pRCC type I, II, and IIM (Dataset S2, Figure S1A-C).

**Figure 1.**
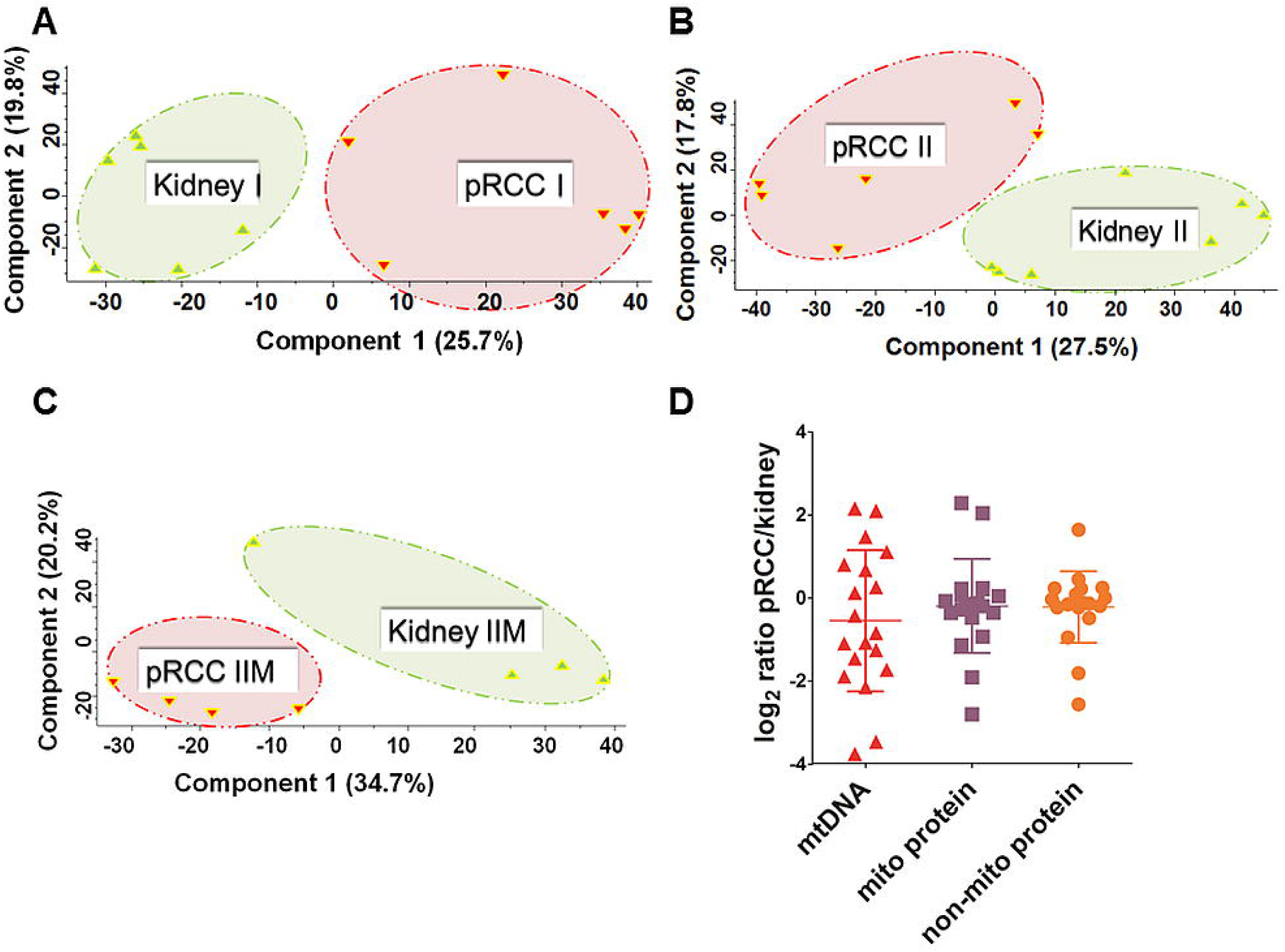
Principal component analysis and evaluation of the mtDNA and mitochondrial protein content in pRCC. PCA analysis in (A) pRCC type I (n=6), (B) pRCC type II (n=6), and (C) pRCC type IIM (n=4) revealed spatial separation for the proteome profiles. (D) Log_2_ ratios of the mtDNA (mean −0.55 ± 1.7 SD) based on WES read depths, the mitochondrial- (mean −0.19 ± 1.13 SD) and non-mitochondrial proteome (−0.22 ± 0.86 SD) in pRCC versus controls, (n=19).

**Table 1.** Clinical and pathologic features of the pRCC cohort.

| Case ID | Age at surgery | Gender | Pathologic T grade | Fuhrman grade | Tumor size (cm) |
| --- | --- | --- | --- | --- | --- |
| C1 type I | 68 | Male | pT3a | 3 | 5 |
| C2 type I | 83 | Male | pT1b | 1 | 6.5 |
| C3 type I | 60 | Male | pT1a | 2 | 3.9 |
| C4 type I | 74 | Male | pT1b | 2 | 6 |
| C5 type I | 35 | Male | pT3a | 2 | 3.5 |
| C6 type I | 49 | Male | pT1b | 2 | 4.2 |
| C7 type I | 59 | Male | pT1b | 2 | 4.5 |
| C1 type II | 54 | Male | pT3b | 3 | 16 |
| C2 type II | 70 | Male | pT1a | 2 | 2.5 |
| C3 type II | 57 | Male | pT2b | 3 | 12.5 |
| C4 type II | 80 | Female | pT1b | 2 | 6.2 |
| C5 type II | 77 | Male | pT3b | 2 | 6 |
| C6 type II | 81 | Female | pT3a | 3 | 8.5 |
| C7 type II | 71 | Male | pT2a | 3 | 7.5 |
| C1 type IIM | 42 | Male | pT3a | 3 | 6.5 |
| C2 type IIM | 72 | Male | pT3a | 3 | 17 |
| C3 type IIM | 67 | Female | pT3a | 3 | 5.5 |
| C4 type IIM | 80 | Male | pT2b | 3 | 11 |
| C5 type IIM | 32 | Male | pT3a | 3 | 10.3 |

### mtDNA Mutations in pRCC did not Reveal any Major Impact on the Respiratory Chain

The assembly of mitochondrial whole exome sequencing (WES) reads derived from 19 patients with pRCC and matched with adjacent healthy kidney tissues showed adequate coverage and quality for reliable mtDNA reconstruction and variant calling (Dataset S3). Mitochondrial mean coverage read depth and mitochondrial assembled bases in the WES dataset ranged from 12.42X to 371.41X and from 91.21% to 100%, respectively. The mtDNA content was 46% reduced (Figure 1D), based on log_2_ ratios of the mtDNA WES read depths between pRCC and matching controls, which is in line with a previous observation in RCC by Southern blot (Meierhofer, Mayr et al. 2004). Furthermore, the abundance of proteins located within the mitochondrion versus all non-mitochondrial proteins showed no difference (Figure 1D).

A total of 260 somatic mtDNA mutations were detected in pRCC samples. Altogether 86 mutations were located within the protein-coding genes, divided into 44 synonymous, 40 non-synonymous and two nonsense mutations. Among the non-synonymous variants, 25 showed a disease score higher than the threshold (>0.7) and a nucleotide variability that was lower than the nucleotide variability cutoff (0.0026). A total of 197 germline mutations were detected, but only one of the seven non-synonymous germline variants were shown to be potentially pathogenic (Dataset S4).

Although 25 somatic non-synonymous and potentially pathogenic events were identified, it was not possible to infer a strong relationship with pRCC, considering that all mutations were found in only ten of the 19 tumors. None of the somatic mutations were shared between the different subjects and homoplasmic rates were generally very low (Dataset S4), however the number of studied cases was too low to draw any final conclusions.

An analysis of copy number variations (CNV) revealed a fragmented pattern of chromosomal gains and losses spread over all chromosomes in all pRCC types (Figure S2), but no clear chromosomal patterns were identified.

### Significantly Decreased Enzymatic Activity of the Respiratory Chain in pRCC

A gene set enrichment analysis (GSEA) was conducted to identify significantly rewired metabolic pathways in the tumor. Significant decreases in all three investigated pRCC types were found in the Kyoto Encyclopedia of Genes and Genomes (KEGG) pathways for: oxidative phosphorylation, the TCA cycle, branched chain amino acids, cytochrome P450 drug metabolism, peroxisomes, fatty acid metabolism, and several amino acid metabolism pathways. (Figure 2A, Dataset S5). The OXPHOS system was the most severely reduced in all pRCC types. Interestingly, there was no obviously different regulation between the three types of pRCC that was detectable at the protein pathway level. The three most significantly increased KEGG pathways in all pRCCs were the spliceosome, the ribosome and the cell cycle (Figure 2A; Dataset S5). An aberrantly increased rate of ribosome biogenesis has been recognized as a hallmark of many cancers, caused by hyperactivation of RNA polymerase I transcription and ribosome biogenesis factors, reviewed in (15, 16).

**Figure 2.**
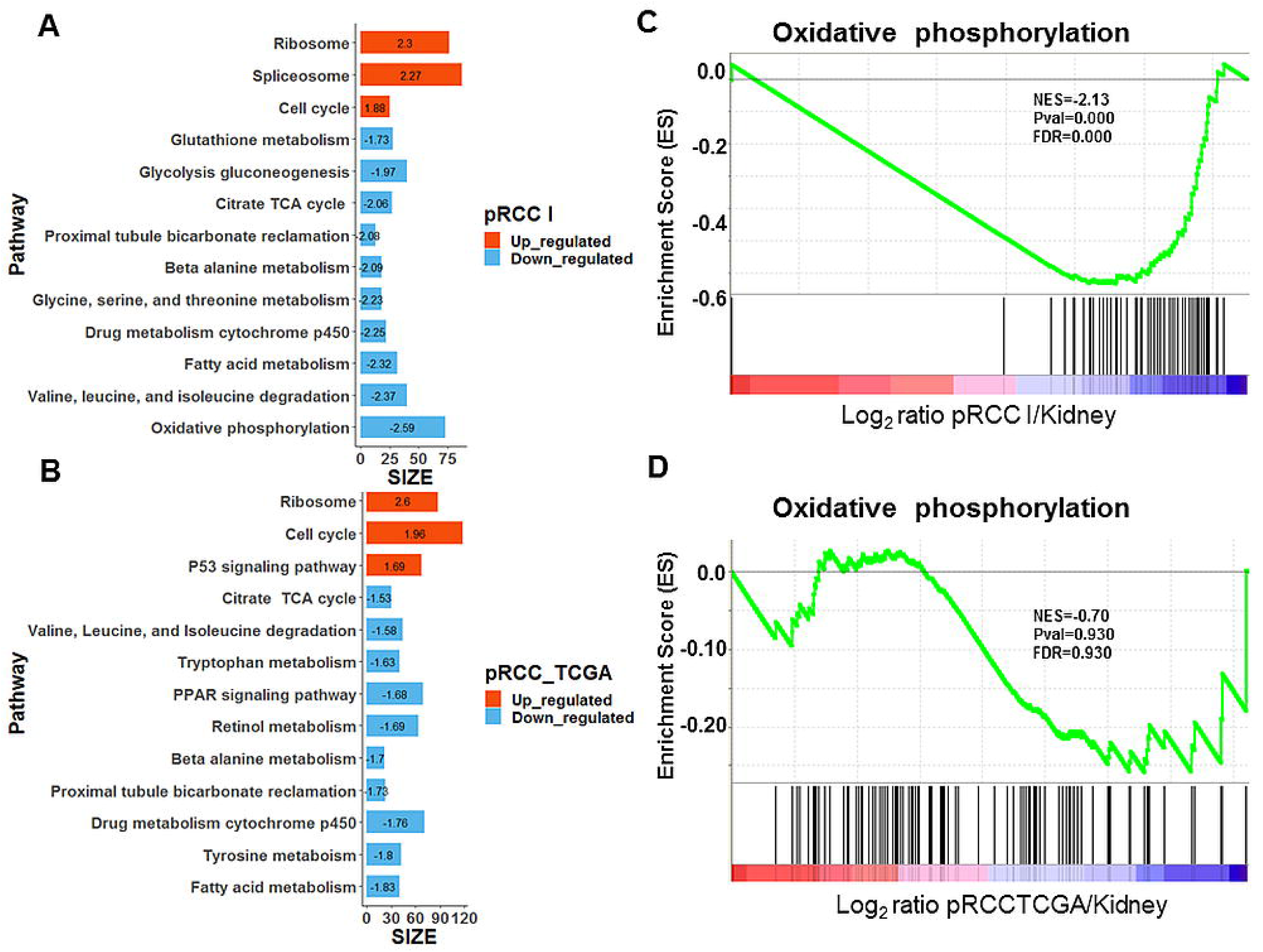
Significantly regulated KEGG pathways between the proteomes of (A) pRCC type I and healthy kidneys and (B) of the transcriptome data retrieved from TCGA. Shown is a collapsed list, the applied cutoff is p ≤ 0.05 and q ≤ 0.1. Specific enrichment plots for the KEGG pathway “oxidative phosphorylation” are shown for (C) the proteome and (D) the transcript data from TCGA. V-ATPases were removed from the pathway “oxidative phosphorylation” displayed in C and D, as they are wrongly assigned in this KEGG pathway. Normalized enrichment score, NES; Size, number of proteins/genes identified within a pathway.

In order to investigate the regulation of the respiratory chain between pRCC and controls in more detail, separate gene sets for all five OXPHOS complexes were created, including the assembly factors. This revealed a reduction in protein abundance for all complexes with the highest observed for complex I (CI) in pRCC, exemplarily shown for type I tumors (Figure 3A). Only assembly factors, that weren’t part of the final complexes, were not decreased.

**Figure 3.**
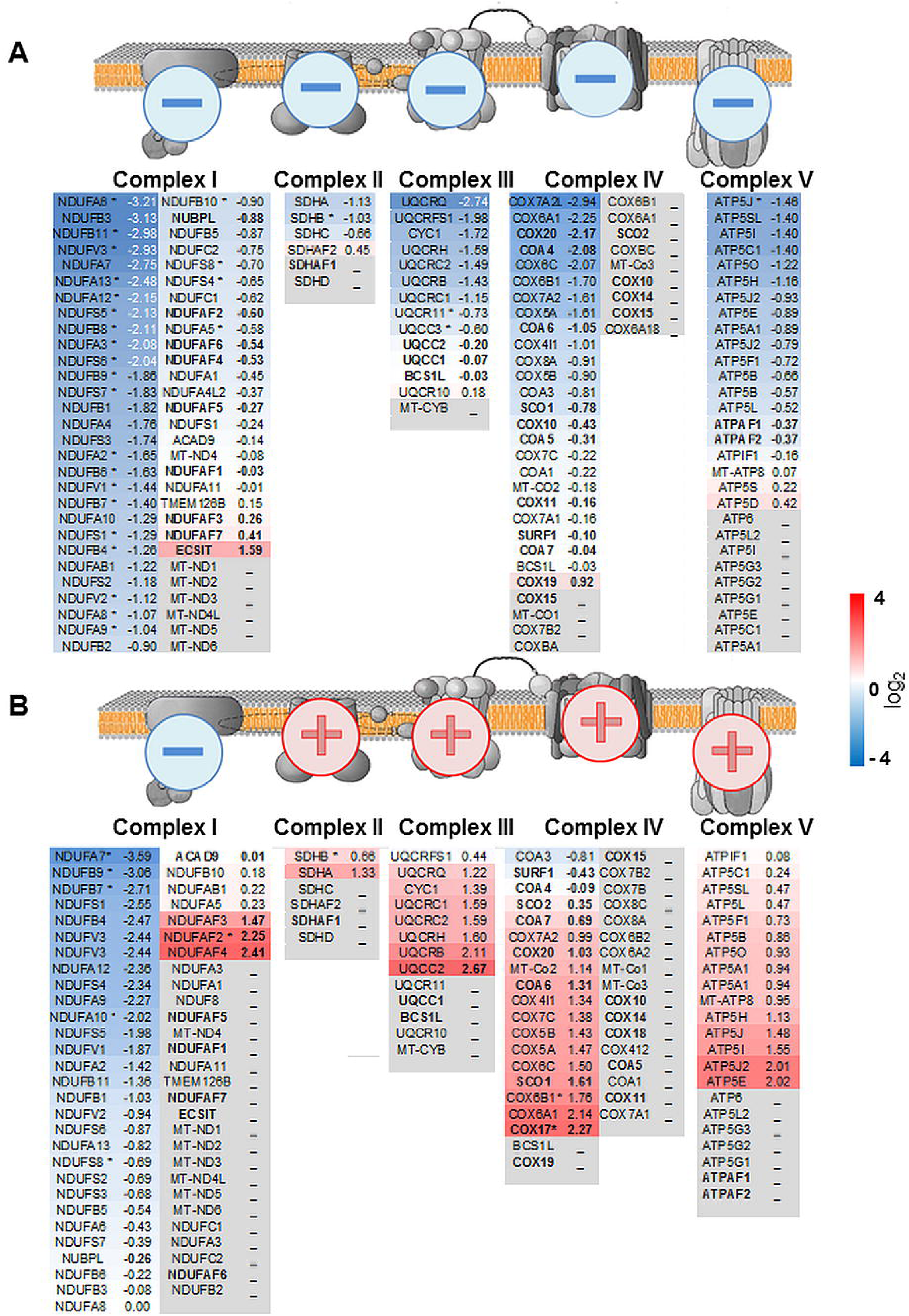
Protein abundance ratios for all individual complexes of the respiratory chain, shown for (A) pRCC type I and (B) renal oncocytomas. Illustrated are schemes of the four OXPHOS complexes and the F_0_F_1_ATPase, including subunits and assembly factors and the according log_2_ fold change between pRCC or renal oncocytomas versus kidney samples. The color gradient intensity in the subunit expresses the low (blue) or high (red) abundance of this protein in the tumor. Assembly factors, not part of the final complexes are shown in bold. * indicates significantly regulated proteins.

### Anti-correlation of Transcripts and Proteins of the Respiratory Chain in pRCC

Altogether, 291 existing pRCC and 32 control transcriptome data retrieved from TCGA (ID: KIRP) (5) were used to clarify the correlation between the abundance of proteins and the expression of transcripts in pRCC versus controls (Dataset S6). GSEA was performed and identified that most of the significantly regulated pathways (Figure 2B, Dataset S7) were very similarly regulated for our proteome study. The ribosome and cell cycle were significantly increased in both omics datasets, whereas the TCA cycle, drug metabolism, fatty acid metabolism, and the pathways involved in amino acid metabolism were all significantly decreased (Figure 2A-B; Dataset S5 and S7). The only striking differences were the “spliceosome” pathway, which was significantly up-regulated only in the pRCC proteomes and the “oxidative phosphorylation” pathway, which was the most decreased pathway at the protein level (Figure 2C), but that was unchanged at the transcript level (Figure 2D). Since this KEGG pathway also includes V-ATPases, which are not part of the respiratory chain, we manually removed them from the analysis. The enzymatic activities of the individual respiratory chain complexes and citrate synthase (CS) were measured for pRCC and the adjacent matching healthy tissues. This revealed a significant reduction in all enzymatic activities of the respiratory chain, the F_0_F_1_ ATPase, and CS in pRCC (Figure 4A-F). Thus, the regulation of the respiratory chain was not determined by the abundance of the respective transcripts but was correlated with the abundance and enzymatic activity of the OXPHOS complexes.

**Figure 4.**
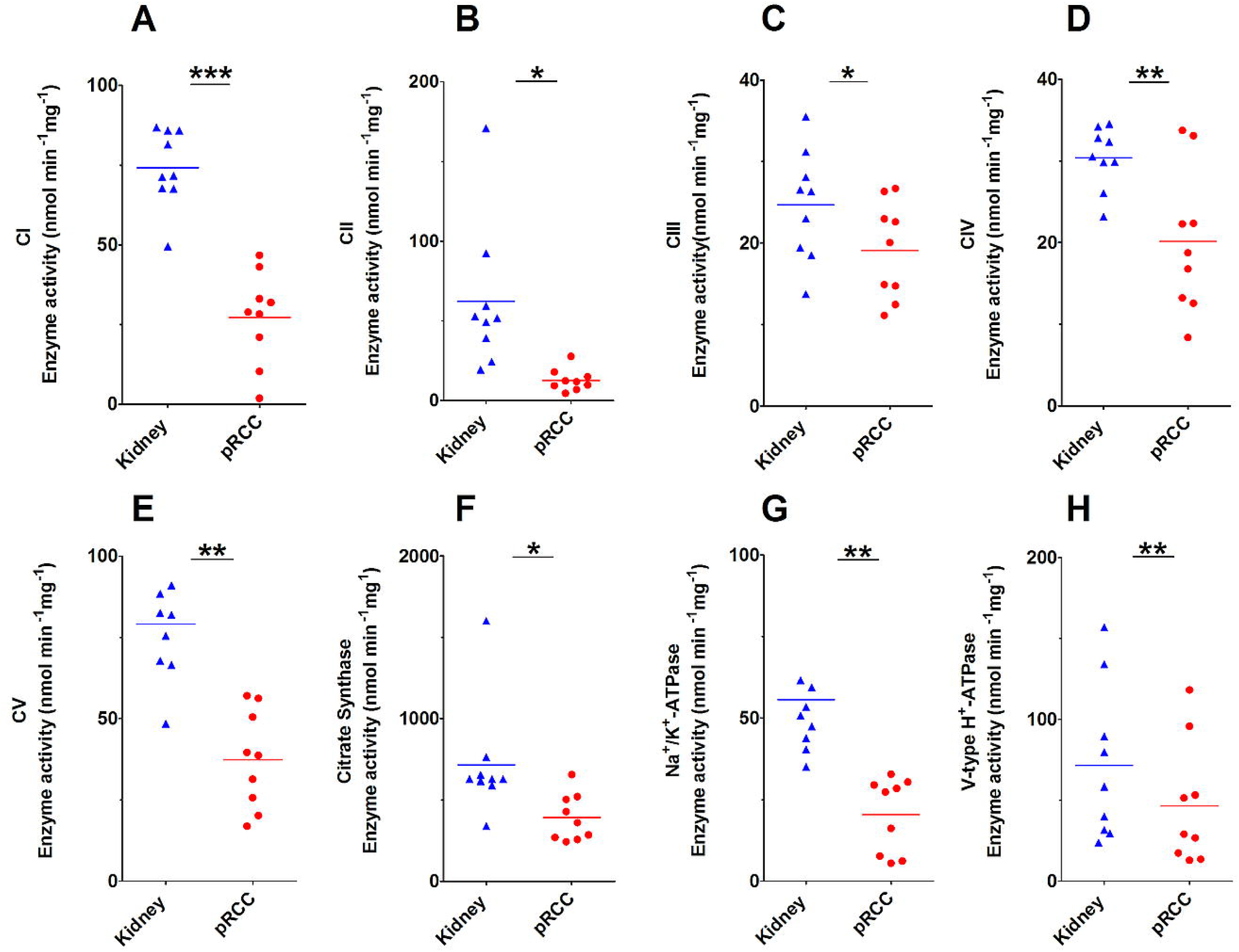
Enzymatic activities of the respiratory chain, F_0_F_1_ ATPases, citrate synthase, and the P- and V-ATPases in pRCC versus kidney control tissues. (A) Complex I (CI), (B) Complex II (CII), (C) Complex III (CIII), (D) Complex IV (CIV), (E) F_0_F_1_ ATPase (CV), (F) Citrate synthase (CS), (G) P-ATPases, (H) V-ATPases. P-values are: *P < 0.05, **P < 0.01, and ***P < 0.001 by a paired t-test; nmol min/mg protein. n= 3 for each type I, II, and IIM.

### Comparison of the Abundance of Proteins Involved in the Respiratory Chain between Malignant pRCC and Benign Renal Oncocytomas

In our previous study on renal oncocytomas, we identified a coordinated up-regulation of proteins of the OXPHOS complexes II-V and the mtDNA, but a striking reduction in the abundance of CI proteins (Figure 3A) (17). This was explained by several specific low-level heteroplasmic mtDNA mutations of the CI genes in renal oncocytomas. In contrast, all pRCC tumor types featured a general reduction in all OXPHOS complexes (Figure 3B) where mtDNA mutations seemed to play no major role, but where the number of mtDNA molecules correlated with respiratory chain protein abundances between these tumor entities.

### pRCCs have Significantly Decreased Levels of V- and P-ATPases

Vacuolar-type H^+^-ATPases (V-ATPase) acidify a wide array of intracellular organelles and pump protons via ATP hydrolysis across intracellular and plasma membranes. Acidity is one of the main features of tumors, V-ATPases control their microenvironment by proton extrusion into the extracellular medium (18). This allows secreted lysosomal enzymes to work more efficiently to degrade the extracellular matrix and promote cellular invasion. In contrast to almost all other cancer types, the V-ATPases were found to be down-regulated in pRCC: This specifically applies to the V-type proton ATPase subunit B, kidney isoform (ATP6V1B1), 21-, 14-, 13-fold (fold changes are shown sequentially for type I, II, IIM, respectively) and others such as ATP6V1H, 3-, 4-, 4-fold; ATP6F1, 3-, 3-, 2-fold; ATP6V1E1, 3-, 3-, 4-fold; and ATP6V1A, 2-, 2-, 2-fold; (Dataset S2). This seems to be a specific feature of pRCC and might play a key role in malignancy.

Similar to the V-ATPases, P-type cation transport ATPases, and the subfamily of Na^+^/K^+^ ATPases were decreased in pRCC (ATP1A1, 4-, 4-, 5-fold; ATP1B1, 4-, 6-, 5-fold; Dataset S2). These ATPases are an integral part of the membrane proteins responsible for establishing and maintaining the electrochemical gradients of Na^+^ and K^+^ ions across the plasma membrane. They are also important for osmoregulation, sodium-coupled transport of several organic and inorganic molecules, and the electrical excitability of nerve and muscle. The enzymatic activities of V- and P-ATPase types were evaluated and showed a significantly reduced activity in pRCC tissues (Figure 4G-H), which correlated to the observed protein abundances.

### Anabolic Glucose Synthesis was Abated in pRCC

The KEGG pathway “glycolysis and gluconeogenesis” was significantly reduced in pRCC (Dataset S5). This pathway describes two opposing functions: the metabolic generation of pyruvate from glucose and the anabolic synthesis of glucose from different substrates, such as glycerol, lactate, pyruvate, propionate, and gluconeogenic amino acids. A more detailed view of each metabolic pathway showed that the abundance of all glycolytic enzymes was either unchanged or increased, whereas all enzymes solely involved in gluconeogenesis were significantly reduced in most pRCCs, such as for PC (17-, 16-, 7-fold), PCK1 (124-, 136-, 388-fold), PCK2 (12-, 11-, 45-fold), ALDOB (100-, 208-, 145-fold), FBP1 (5-, 12-, 49-fold), and FBP2 (not detected (nd), nd, 53-fold) (Figure 5A-C, Dataset S2). Interestingly, the two fructose-bisphosphate aldolase isoforms A (2-, 2-, 5-fold) and C (1-, 1-, 4-fold) were instead increased or unchanged in pRCC (Figure 5D). These isoforms have a high affinity for fructose-bisphosphate (FDP) to foster glycolysis, whereas the highly diminished isoform B has a low affinity for FDP and hence converts the back-reaction from glyceraldehyde-3-phosphate to FDP during gluconeogenesis (19, 20). A similar decrease in the respective transcripts was found in the TCGA data, where *ALDOB* was 600-fold and *ALDOC* 2-fold decreased and *ALDOA* 2-fold increased. A specific and regulated interaction between ALDOB and the rate-limiting gluconeogenic enzyme fructose-1, 6-bisphosphatase 1 (FBP1) has been shown. This result confirms the view that ALDOA and ALDOB play different roles in glucose metabolism (21). The shut-down of the gluconeogenic pathway was thus one of the most relevant metabolic changes observed and can be regarded as a metabolic hallmark in pRCC.

**Figure 5.**
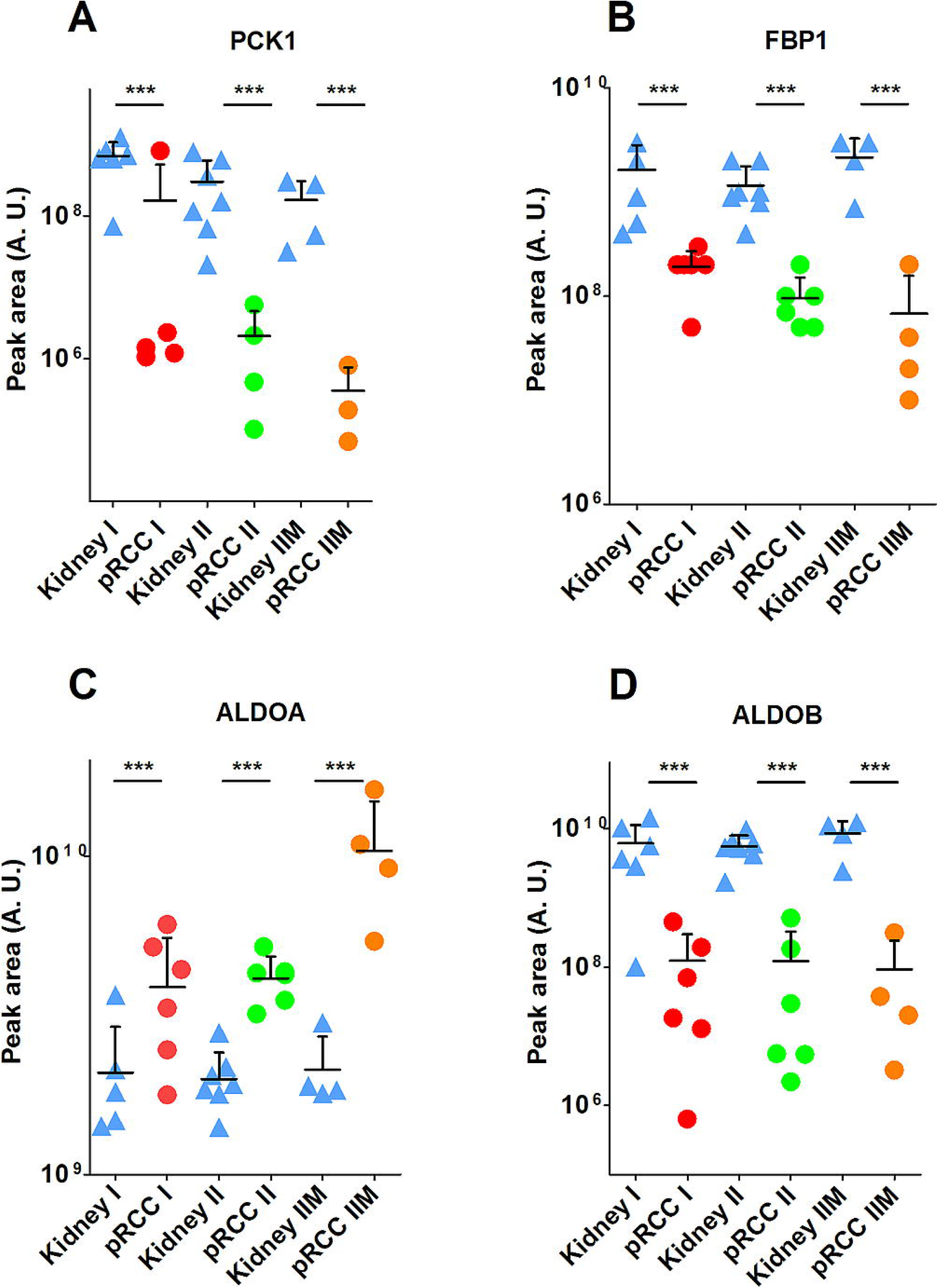
Protein abundances of gluconeogenic enzymes in pRCC and normal kidney tissues. (A) PCK1, phosphoenolpyruvate carboxykinase 1; (B) FBP1, fructose-1,6-bisphosphatase; (C) ALDOA, fructose-bisphosphate aldolase A; (D) and ALDOB. P-value is: ***P < 0.001 by a two-tailed student’s t-test.

### Dramatically Increased Glutathione Levels in pRCC are Based on Glutamine Consumption

Metabolome profiling revealed a highly significant increase in reduced glutathione (GSH, 47-, 68-, 219-fold; fold changes are shown sequentially for type I, II, IIM, respectively) and oxidized glutathione (GSSG, 871-, 847-, 6,707-fold) in pRCC types, Figure 6A-F, Dataset S8). A case by case specific GSH/GSSG ratio was calculated (Figure 6F) and revealed that there was a 10-fold average increase in oxidative stress burden in the tumor.

**Figure 6.**
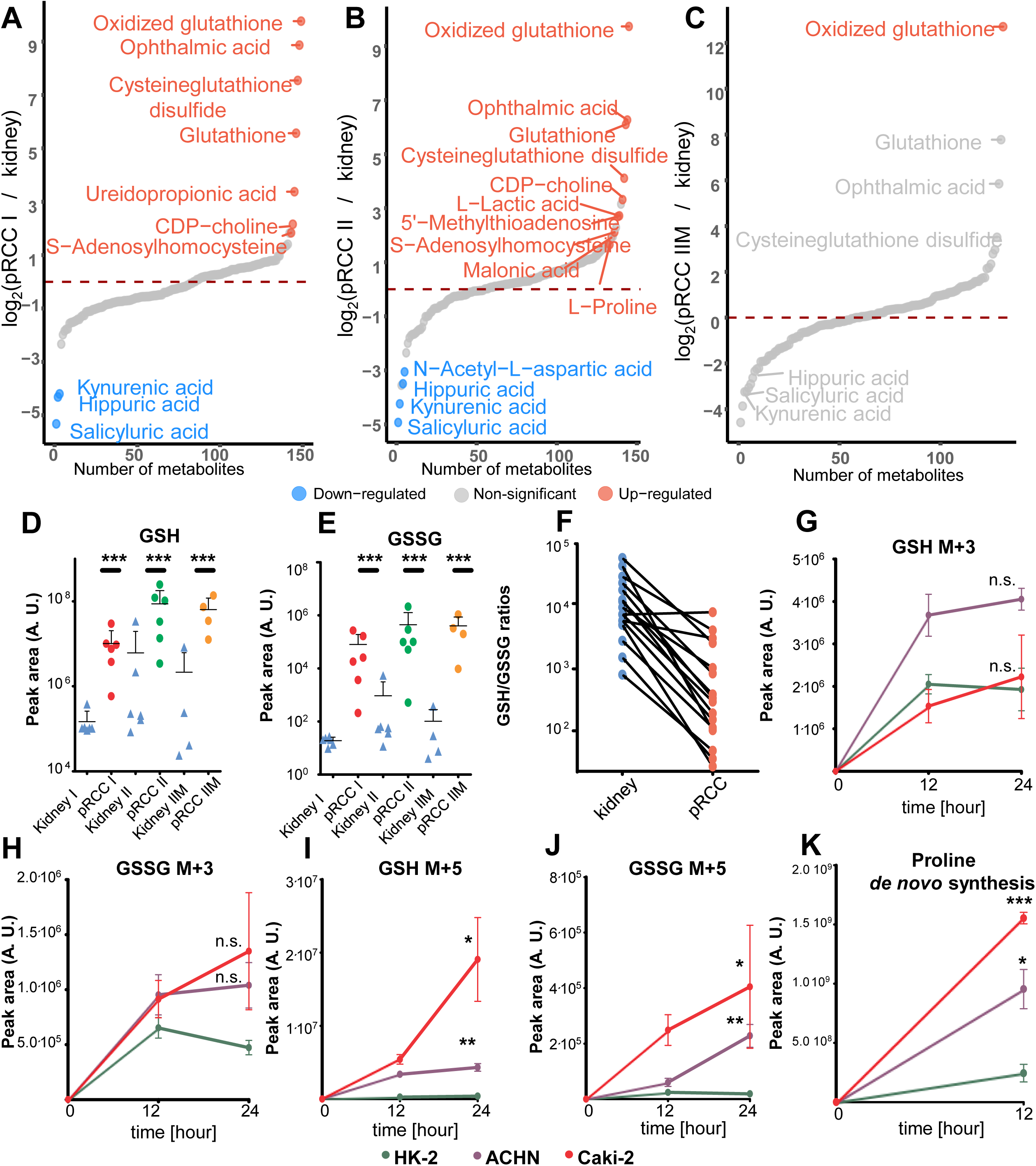
Metabolome profiling, specific abundancies of reduced and oxidized glutathione in tissues, and *de novo* synthesis of glutathione in pRCC. Metabolome profile of (A) pRCC type I (n= 6), (B) pRCC type II (n= 6), and (C) pRCC type IIM (n= 4) versus kidney tissues. Significantly (FDR < 0.05) up- and down-regulated metabolites are shown in red and blue, respectively. (D) Relative abundance of glutathione (GSH) and (E) of oxidized glutathione (GSSG) in the tissues and (F) the according case-specific GSH/GSSG ratios. Metabolic tracing of (G-H) ^13^C_6_ glucose and of (I,J) ^13^C_5_^15^N glutamic acid to monitor GSH (G,I) *de novo* synthesis and its oxidation product GSSG (H,J) in the pRCC cell lines Caki-2 and ACHN compared to HK-2 control cells. (K) Proline *de novo* synthesis based on ^13^C_5_^15^N glutamic acid in the same cell lines. Mean ± SD, n=3 cell culture replicates, P-values: *P < 0.05, **P < 0.01, and ***P < 0.001 by a paired t-test.

Glutathione functions as a cellular redox buffer for detoxification and can be either synthesized *de-novo* or imported via the glutathione salvage pathway, where extracellular GSH is cleaved by γ-glutamyltranspeptidases (GGTs) (22). Furthermore, ophthalmic acid, a tripeptide analog of glutathione, was increased (468-, 77-, 58-fold) in pRCC. It was described as a byproduct of glutathione synthetase (GS) and γ-glutamylcysteine synthetase (GCS) and as a new biomarker of oxidative stress (23).

Remarkably, even though metabolites involved in glutathione metabolism were significantly increased in pRCC, the abundance of glutathione synthetase (GSS) was unchanged (1-, 1-, 1; Dataset S2). Glutamate cysteine ligase (GCL) is the rate-limiting step in GSH biosynthesis. Only the pRCC type IIM had significantly elevated levels of GCLM (1-, 2-, 10-fold), the regulatory subunit of this enzyme alleviates the feedback inhibition of GSH together with the catalytic subunit GCLC (24).

Cysteineglutathione disulfide, which can react with protein thiol groups, causing the reversible post-translational modification S-glutathionylation for transducing oxidant signals (25) was (187-, 17-, 11-fold) elevated. In contrast, the glutathione S-transferase A2 (GSTA2, 146-, 36-, 85-fold), microsomal glutathione S-transferase 1 (MGST1, 5-, 1-, nd-fold), and glutathione S-transferase Mu 2 and 3 (GSTM2, 5-, 5-, 3-fold; GSTM3, 8-, 10-, 5-fold), which conjugate reduced glutathione to a wide number of exogenous and endogenous hydrophobic electrophiles, were reduced in pRCC, but other enzymes were unchanged, such as GSTT1, GSTP1, GSTK1, and GSTO1. Glutathione peroxidase 3, which protects cells and enzymes from oxidative damage by catalyzing the reduction of hydrogen peroxide, lipid peroxides, and organic hydroperoxide was also significantly reduced in pRCC (GPX3, 6-, 5-, 6-fold).

The two identified glutathione transferases, which catalyze the conjugation of GSH to xenobiotic substrates for detoxification, were also strongly reduced in pRCC, e.g. glutathione hydrolase 1 proenzyme GGT1 (3-, 4-, 4-fold) and GGT5 (20-, 8-, 17-fold). They cleave the gamma-glutamyl peptide bond of glutathione conjugates, the only one identified was gamma-glutamyl lysine, which was increased (2-, 3-, 3-fold) in pRCC. Gamma-glutamyl amino acids can be further metabolized by γ-glutamyl cyclotransferase (GGACT) (26), whose level was decreased (13-, 45-, 108-fold) to produce pyroglutamic acid (5-oxo-proline), which was not significantly regulated (2-, 2-, −2-fold) as well as other amino acids.

These data show a notable rewiring of the entire process of glutathione metabolism in pRCC, however metabolite and protein abundances do not necessarily reflect the metabolic flux. To address the question, as to whether GSH is continuously made by *de novo* synthesis and which substrates significantly contribute to its synthesis, two isotope tracing experiments were performed in the pRCC derived cell lines Caki-2 and ACHN and the kidney control cell line HK-2 at time points 0, 12, and 24 hours. The first experiment employed ^13^C_6_ labeled glucose, the second ^13^C_5_ N glutamic acid as a tracer, a generalized labeling scheme for both is depicted in figure 7.

**Figure 7.**
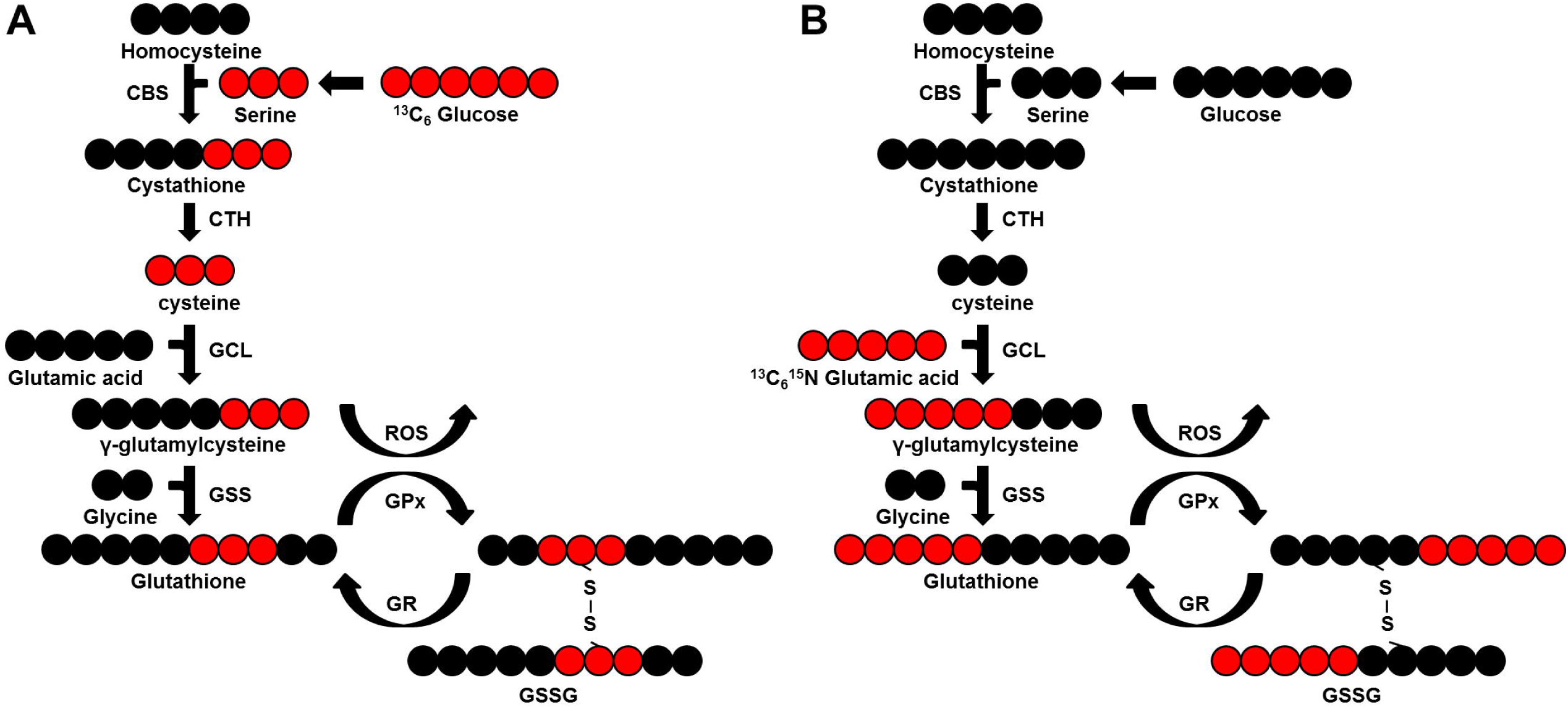
Scheme of the *de novo* synthesis pathway of glutathione. (A) based on ^13^C_6_ glucose and (B) on ^13^C_5_^15^N glutamic acid as tracers. Red circles, ^13^C_6_; black circles, ^12^C_6_. CBS, cystathionine b-synthetase; CTH, cystathionase; GCL, glutamate-cysteine ligase; GSS, glutathione synthetase; GPx, glutathione peroxidase; GR, glutathione reductase; GSSG, oxidized GSH. Of note, GSSG can exist in three versions in parallel, as it is made of two GSH molecules: ^13^C_6_, ^13^C_3_, and ^12^C when ^13^C_6_ glucose is the tracer and ^13^C_10_^15^N_2_, ^13^C_5_^15^N, and ^12^C^14^N when ^13^C_5_^15^N glutamic acid is the tracer.

Only a slight increase was observed for the *de novo* synthesis of GSH and GSSG in Caki-2 cells (1-fold-, 2.8-fold at 24h) and ACHN cells (2.1-fold-, 2.2-fold at 24h) compared with HK-2 cells when ^13^C_6_ glucose was used as the probe (Figure 6G,H). In contrast to this, a significant and dramatic increase of GSH and GSSG *de novo* synthesis was observed for both, Caki-2 (48.5-fold-, 10.8-fold at 24h) and ACHN (11-fold-, 12-fold at 24h) cells, when ^13^C_5_ N glutamic acid was used as a probe (Figure 6I,J). Our data demonstrated that GSH *de novo* synthesis is significantly increased in pRCC cell lines and is based on glutamine consumption, a precursor of glutamate.

### Glutamine is the Main Nutrient Source in pRCC

In general, glucose and glutamine are the main carbon sources in eukaryotic cells. Tumor cells frequently reduce OXPHOS capacity and are therefore even more dependent on nutrient consumption. Proteome profiling showed that the abundance of proteins involved in proline synthesis, such as CAD protein (CAD, 2-, 5-, 10-fold), ALDH18A1, 1-, 2-, 14-fold; PYCR1, nd, nd, 115-fold; and PYCR2, 5-, 7-, 15-fold, Dataset S2) increase from type I over type II leading to the highest values in metastatic type IIM, indicating a metabolic shift towards proline *de novo* synthesis originating from glutamine. Based on these results, an isotope tracing experiment was performed in pRCC derived cell lines Caki-2 and ACHN versus HK-2 kidney controls to quantify the consumption of isotopically labeled glutamate for proline *de novo* synthesis.

Tracing of ^13^C_5_ N glutamic acid revealed a significant and (6.3-, 3.9-fold) increase of C_5_ N proline after 12 hours in the pRCC derived cell lines Caki-2 and ACHN versus HK-2 (Figure 6K). The high flux in pRCC cell lines correlates well with the abundance of protein in pRCC tissues of enzymes involved in this pathway showing that these tumor cell lines are dependent on glutamine consumption.

## Discussion

pRCCs are well characterized at the genomic level, with several driver mutations and chromosomal rearrangements having been identified (5). How these alterations translate to proteome- and metabolome regulation are not well understood, but they determine the fate and progression of tumors. Multi-omics profiling of pRCC was performed, revealing a fundamental reprogramming of the pathways for gluconeogenesis, the respiratory chain, and for glutathione metabolism. These can be regarded as a general hallmark of kidney tumors, as was previously observed in renal oncocytomas (17), chRCCs (27), and in ccRCCs at the transcript level (28).

The anti-correlations that have been identified between genetic and non-genetic profiling argue for focusing on these so far under-studied fields.

Gluconeogenesis, an anabolic and highly endergonic pathway, generates glucose from small carbohydrate precursors, for example from lactate during intense exercise, or over periods of fasting and starvation. This pathway is also regarded as an essential process for tumor cell growth (29), since biosynthetic reactions in cancer cells are highly dependent on glycolytic intermediates (30). The kidney may be nearly as important as the liver in gluconeogenesis (31) and pRCCs have been shown to moderately accumulated glucose, which further increases the already higher Fuhrman grades (32). This might be the reason why pRCCs reduce this endergonic pathway, since enough glucose can be imported. Blocking of mTOR activity was shown to augment shuttling of pyruvate into gluconeogenesis, which results in futile cycling of glucose that finally leads to a halt in cancer cell proliferation and ultimately to cell death (33). Equal amounts of amino acid levels were detected in pRCC and kidney tissues in our study, supporting the idea of a sufficient nutrient supply.

The gluconeogenic gene *FBP1* was previously found to be down-regulated in over 600 ccRCCs and was associated with a poor disease prognosis. Thus, FBP1 has been shown to fulfill two distinct functions, by antagonizing glycolytic flux and thus inhibiting the Warburg effect, and by inhibiting the nuclear function of HIF in a catalytic-activity-independent manner, leading to reduced expression of HIF targets such as VEGF, LDHA, and GLUT1 (28). This unique dual function of the FBP1 protein explains its ubiquitous loss in ccRCC, distinguishing it from other tumor suppressors that are not consistently mutated in all tumors (28).

Moreover, the inhibition of FBP1 leads to the activation of AMP-activated protein kinase (AMPK). The aldolases (A-C) are required for the formation of a super lysosomal complex containing V-ATPase, ragulator, axin, liver kinase B1 (LKB1), and AMPK in its active form (34). AMPK activation plays a central role in glucose sensing at the lysosome and acts contrary to other regulatory systems such as the mammalian target of rapamycin (mTOR).

A general shut-down of the entire gluconeogenesis pathway in pRCC was also identified at the proteome level in our study, which has fundamental implications for the metabolic regulation of a cell and organ. Specifically, the two aldolase isoforms A and C, which foster glycolysis were unchanged, but the aldolase isoform B, necessary for gluconeogenesis was greatly diminished in pRCC. This again indicates a critical metabolic change, as previously observed in chRCC (27) and in gastric cancer (35). Conversely, an over-expression of gluconeogenic genes and proteins is frequently found in other tumor species, such as the over-expression of *ALDOB* in ccRCC (36) and colon cancer (37–39), which was also associated with tumor progression and poor prognosis.

Also the up-regulation of *ALDOA* has been reported in many cancer types, such as oral squamous cell carcinoma (40), osteosarcoma (41), lung cancer (42), and hepatocellular carcinoma (43). Specifically, an increase in ALDOA was shown for all RCC types and was associated with metastasis, histological differentiation, and poor prognosis. Furthermore, silencing ALDOA expression in ccRCC cell lines decreased their proliferative, migratory, and invasive abilities, while ALDOA overexpression increased these abilities (44).

In addition, the abundance and enzymatic activities of the P- and V-ATPases were found to be significantly reduced in our pRCC panel. A possible mechanism by which V-ATPases are thought to contribute to cancer cell migration and invasion is to acidify extracellular space to promote the activity of acid-dependent proteases that are involved in invasion (18, 45, 46). Besides the classical role of regulating acidity within a cell, recent studies showed that V-ATPases, as part of the V-ATPase-Ragulator complex, serve as a dual sensor for energy/nutrient sufficiency and deficiency and they can initiate the metabolic switch between catalytic and anabolic pathways (47). It has been further shown and that glycolysis is directly coupled to the V-ATPases by protein-protein interactions (48, 49).

The significant reduction of enzymes involved in gluconeogenesis thus has metabolic consequences on multiple layers and is a hallmark of all investigated kidney cancers, such as in pRCC (this study), ccRCC (28), chRCC (27), and renal oncocytomas (17). By abandoning this endergonic pathway in pRCC the tumor is able to simultaneously reduce other processes involved in the generation of ATP. Indeed, pathways involved in fatty acid metabolism, amino acid metabolism, as well as OXPHOS and the TCA cycle pathways were significantly down-regulated in our pRCC specimen.

Another frequently observed phenomenon in cancer is the diminished oxidative phosphorylation capacity, known already for decades as the “Warburg effect”. The abundance of all proteins involved in oxidative phosphorylation and the F_0_F_1_ATPase as well as the corresponding enzymatic activities were significantly reduced in our pRCC panel. This was previously shown for chRCC (27) and only for OXPHOS enzymatic activities and the mtDNA content in ccRCC and pRCC (50). In contrast, the enhanced expression of LDH-A and lactic acid was observed in our study and this has been associated with aggressive and metastatic cancers in a variety of tumor types (51–53).

By comparing our proteome-with transcriptome data from TCGA (5), the main differentially regulated pathway was found to be the respiratory chain, which was the most highly decreased pathway on the protein level, but unchanged at the transcript level. A similar discrepancy between transcripts and proteins in the regulation of the respiratory chain was previously observed by us in benign renal oncocytomas (17) and malignant chRCC (27). Enzymatic activities of the respiratory chain in pRCC and in renal oncocytomas (54) and chRCC (27) matched with protein abundances rather than gene expression. The mechanism for this anti-correlation still remains elusive, but might be directly correlated to the decreased mtDNA level, or also caused by the interference of miRNAs and the stability of transcripts or proteins. This demonstrates the necessity of surveying multiple omics profiles.

The most strikingly increased set of metabolites in pRCC were those involved in glutathione metabolism (GSH, GSSG, cysteine-glutathione disulfide, ophthalmic acid). This is similar to those previously identified in renal oncocytomas (17, 55) and chRCC (27, 56). GSH is an important ROS scavenger (57) and frequently produced by several tumor types to withstand unusual levels of oxidative stress (22). Therefore increased GSH levels in pRCC may be considered as the main strategy for the tumor to overcome ROS stress originating from a dysregulated respiratory chain.

By probing the metabolic flux for GSH synthesis, a significant increase of the synthesis rate was observed in pRCC derived cell lines over kidney controls when using glutamate as a substrate. This is in agreement with another study, which found that glutamine dependence in ccRCC suppresses oxidative stress (58). The inhibition of GSH synthesis by a specific glutaminase (GLS) inhibitor and the simultaneous treatment with hydrogen peroxide resulted in a high apoptosis rate in ccRCC (58). Hence, additional administrationof antioxidants during (chemotherapeutic) cancer treatment have been frequently shown to have no or even pro-tumor effects (59, 60). High GSH levels in RCC, which protect the tumor from increased ROS stress, should therefore be therapeutically exploited by reducing the antioxidant levels, of GSH, and increasing ROS stress at the same time to a level where healthy cells can still survive, but tumorous cells are forced into apoptosis.

## Conclusion

Key metabolic reprogramming processes, such those for gluconeogenesis, the respiratory chain, and glutathione metabolism are not only the main molecular characteristics for papillary RCC, but rather seem to be a general feature for other kidney tumors as well. Specifically, the reinforcement of glutathione metabolism, reflecting the increased burden of oxidative stress, and abandoning endergonic processes may hold key therapeutic implications as a future treatment option.

## Experimental Procedures

### Tissue Dissection and Verification of Papillary RCC

Malignant and non-malignant tissues of 19 nephrectomies performed between 2008 and 2016 at the Department of Urology, Charité – Universitätsmedizin Berlin, were collected in liquid nitrogen immediately after surgery and preserved at −80°C. The clinical characteristics of the tumors are reported in Table 1. From the collected tissue samples, histologic sections were stained with hematoxylin and eosin. The diagnosis of pRCC and the corresponding matched tumor-free kidney tissue was done according to WHO classification criteria. Only cases with a clear diagnosis of pRCC were considered for the study.

### Whole Exome Sequencing (WES)

DNA was isolated from remaining pellets from metabolite extraction using a DNA purification kit following the manufacturer’s protocol for tissues (QIAmp DNA Mini Kit, QIAGEN, Hilden, Germany). In brief, samples were digested by proteinase K at 56°C overnight and RNase A treated at 70°C, before subjecting to exome sequencing.

The library preparation was performed according to Agilent’s SureSelect protocol (SureSelectXT Human All Exon V5, protocol version B4 August 2015) for Illumina paired-end sequencing. In brief, 200 ng of genomic DNA (in 50 μl low TE) were sheared for 6×60 sec on a Covaris™ S2 (duty factor 10%, intensity 5, 200 cycles per burst).

The fragmented DNA (150-200 bp) was purified using AMPure XP beads and subjected to an end-repair reaction. Following another purification step, the DNA was 3’adenylated and furthermore purified. Paired-end adaptors were ligated and the afterward purified library was amplified with 10 amplification cycles. The amplified library was purified, quantified and hybridized to the probe library for exome capture. Captured fragments were purified using streptavidin-coated beads and eluted with 30 μl nuclease-free water. Using Herculase-enzyme, the enriched libraries were amplified and indexed with barcoded primers followed by cleanup and quantification.

The resulting libraries were pooled and subjected to Illumina NextSeq4000 paired-end sequencing (6 libraries/FC; 2 × 150 bp).

Quantification of the SureSelect captured library: Before sequencing, the samples were re-quantified with two methods. First, the size and concentration was checked on the Agilent 2100 Bioanalyzer and in a second step the enrichment efficiency was estimated by qPCR (Applied Biosystems) using a primer set for an enriched exon (fw: ATCCCGGTTGTTCTTCTGTG and rv: TTCTGGCTCTGCTGTAGGAAG) and a primer set in an intron region as a negative control (fw: AGGTTTGCTGAGGAACCTTGA and rv: ACCGAAACATCCTGGCTACAG). In general, the CT-values of target and control fragments differed by 6 to 10, thus confirming a very good enrichment of our target regions.

After diluting the captured libraries to 10 nM, Genome Analyzer single-read flow cells were prepared on the supplied Illumina cluster station and 36 bp single-end reads on the Illumina Genome Analyzer IIx platform were generated following the manufacturer’s protocol. Images from the instrument were processed using the manufacturer’s software to generate FASTQ sequence files.

### Analysis of mtDNA Mutations

The FASTQ files were used as input for the MToolBox pipeline (61) in order to extract mitochondrial DNA sequences and quantify each variant allele heteroplasmy and related confidence interval. The same pipeline allows haplogroup prediction of mtDNA sequences, detection of mismatches, insertions and deletions and the functional annotation of the identified variants. The *in silico* prioritization criteria (62) were used to target the mitochondrial DNA variants of clinical interest. Thus, variants found in the mitochondrial reference sequences (rCRS, RSRS and MHCS), which occurred in non haplogroup-defining sites with a nucleotide variability lower than the nucleotide variability cutoff (0.0026) and a disease score above the disease score threshold of 0.43 for non-synonymous coding for proteins, 0.35 for tRNA, and 0.60, for rRNA variants were prioritized.

### Sample Preparation for Proteomics

About 10 mg frozen tissue per sample was homogenized under denaturing conditions with a FastPrep instrument (three times for 60 s, 6.5 m × s^−1^) in a buffer containing 4% SDS, 0.1 M DTT, 0.1 M Tris pH 7.8, followed by sonication for 5 min, boiled at 95°C for 5 min and precipitated with acetone at −20°C overnight. Lyophilized proteins were dissolved in 6 M guanidiniumchlorid, 10 mM tris(2-carboxyethyl)phosphine, 40 mM chloroacetamide, and 100 mM Tris pH 8.5. Samples were boiled for 5 min at 95 *^◦^*C and sonicated for 15 min in a water sonicator. The lysates were diluted 1:10 with nine times volume of 10% acetonitrile and 25 mM Tris, 8.5 pH, followed by trypsin digestion (1:100) at 37 *^◦^*C overnight. Subsequent, the peptides were purified with C18 columns. For whole proteome profiling, 90 µg of each sample was fractionated by strong cation exchange (SCX) chromatography. Five µg of each SCX fraction was used for proteome profiling.

### LC-MS Instrument Settings for Shotgun Proteome Profiling and Data Analysis

LC-MS/MS was carried out by nanoflow reverse phase liquid chromatography (Dionex Ultimate 3000, Thermo Scientific, Waltham, MA) coupled online to a Q-Exactive HF Orbitrap mass spectrometer (Thermo Scientific, Waltham, MA). Briefly, the LC separation was performed using a PicoFrit analytical column (75 μm ID × 55 cm long, 15 µm Tip ID (New Objectives, Woburn, MA) in-house packed with 3-µm C18 resin (Reprosil-AQ Pur, Dr. Maisch, Ammerbuch-Entringen, Germany). Peptides were eluted using a gradient from 3.8 to 40% solvent B (79.9% acetonitrile, 20% water, 0.1% formic acid) in solvent A (0.1 % formic acid in water) over 120 min at 266 nL per minute flow rate. Nanoelectrospray was generated by applying 3.5 kV. A cycle of one full Fourier transformation scan mass spectrum (300−1750 m/z, resolution of 60,000 at m/z 200, AGC target 1e^6^) was followed by 16 data-dependent MS/MS scans (resolution of 30,000, AGC target 5e^5^) with a normalized collision energy of 27 eV. In order to avoid repeated sequencing of the same peptides, a dynamic exclusion window of 30 sec was used. In addition, only peptide charge states between two to eight were sequenced.

Raw MS data were processed with MaxQuant software (v1.6.0.1) (63) and searched against the human proteome database UniProtKB with 70,941 entries, released in 01/2017. Parameters of MaxQuant database searching were: A false discovery rate (FDR) of 0.01 for proteins and peptides, a minimum peptide length of 7 amino acids, a mass tolerance of 4.5 ppm for precursor and 20 ppm for fragment ions were required. A maximum of two missed cleavages was allowed for the tryptic digest. Cysteine carbamidomethylation was set as fixed modification, while N-terminal acetylation and methionine oxidation were set as variable modifications. MaxQuant processed output files can be found in Dataset S1, showing peptide and protein identification, accession numbers, % sequence coverage of the protein, q-values, and LFQ intensities. Contaminants, as well as proteins identified by site modification and proteins derived from the reversed part of the decoy database, were strictly excluded from further analysis. The mass spectrometry proteomics data have been deposited to the ProteomeXchange Consortium via the Pride partner repository (64) with the dataset identifier PXD013523.

### Metabolite Extraction and Profiling by Targeted LC-MS

About 30 mg of 16 pRCC and healthy kidney tissues, shock-frozen in liquid nitrogen, was used for metabolite profiling. Metabolite extraction and tandem LC-MS measurements were done as we have previously reported (17, 65). In brief, methyl-tert-butyl ester (MTBE), methanol, ammonium acetate, and water were used for metabolite extraction. Subsequent separation was performed on an LC instrument (1290 series UHPLC; Agilent, Santa Clara, CA), online coupled to a triple quadrupole hybrid ion trap mass spectrometer QTrap 6500 (Sciex, Foster City, CA), as reported previously (66). Transition settings for multiple reaction monitoring (MRM) are provided in Dataset S9. The mass spectrometry data have been deposited in the publically available repository PeptideAtlas and can be obtained via http://www.peptideatlas.org/PASS/PASS01368. The metabolite identification was based on three levels: (i) the correct retention time, (ii) up to three MRM’s and (iii) a matching MRM ion ratio of tuned pure metabolites as a reference (66). Relative quantification was performed using MultiQuant software v.2.1.1 (Sciex, Foster City, CA). The integration setting was a peak splitting factor of 0 and all peaks were reviewed manually. Only the average peak area of the first transition was used for calculations. Normalization was done according to used amounts of tissues and subsequently by internal standards, as indicated in Dataset S8.

### Cell Culture Conditions for Glutathione and Proline *de novo* Synthesis

The two pRCC-derived cell lines Caki-2 (ATCC HTB-47, reclassified from ccRCC (67)) and ACHN (ATCC CRL-1611) and the human kidney (HK-2, cortex/proximal tubule, ATCC CRL-2190) cell line were cultivated in Dulbecco’s modified Eagle medium (DMEM, Life Technologies, New York, NY) containing 4.5 g/L glucose, supplemented with 10% fetal bovine serum (FBS, Silantes, Munich, Germany) and 1% penicillin–streptomycin–neomycin (Invitrogen, Carlsbad, CA) at 37 °C in a humidified atmosphere of 5% CO_2_.

To determine the source for GSH *de novo* synthesis and potential differences between the pRCC cell lines Caki-2 and ACHN versus HK-2 kidney controls, two isotope tracing experiments were performed. The first experiment employed ^13^C_6_ labeled glucose, the second ^13^C_5_^15^N glutamic acid as probe, a scheme of GSH synthesis is outlined in figure 7. In addition, proline *de novo* synthesis was monitored simultaneously within the experimental setting of glutamate as a tracer (Figure 6J).

GSH labeling dynamics were probed by sampling at time points 0, 12, and 24 hours, for proline 0 and 12 hours were taken in 6-well plate triplicates for all three cell lines. The cells were rinsed twice with PBS and replenished at time point 0 by either a glucose-free DMEM medium with the addition of 5 mM ^13^C_6_ glucose (Cambridge Isotope Laboratories Inc., Tewksbury, MA), 10% dialyzed FBS and 1% penicillin/streptomycin, or by HBSS solution supplemented with 2 mM ^15^N-glutamic acid (Cambridge Isotope Laboratories), 5 mM glucose, 0.4 mM glycine, 10% dialyzed FBS and 1% penicillin/streptomycin. Before metabolite extraction, cells were washed twice by PBS, ice-cold methanol (−80°C) was added and the cells were scrapped from the plate and metabolites were extracted as described for tissues. The MRM method was extended to also include isotope-transitions for metabolites originating from labeled glucose and glutamate (Dataset S9). As GSSG can be made of either one or two labeled GSH molecules, both versions were measured (^13^C_6_ glucose: M+3 and M+6; ^13^C_5_ N-glutamic acid: M+5 and M+10).

### Experimental Design, Statistical Rationale, and Pathway Analyses

Seven pRCC type I, seven pRCC type II, and five pRCC type II metastatic cancer samples were compared with adjacent matched normal kidney tissues. For quantitative proteome profiling, nanoscale liquid chromatography coupled to high-resolution mass spectrometry (nano-LC-MS/MS) was used to quantify the abundance of dysregulated proteins. For quantitative metabolome profiling, a UHPLC coupled to QTrap instrument was used for the targeted approach (multiple reaction monitoring, MRM) to identify and quantify the abundance of dysregulated metabolites.

For proteome and metabolome data sets, a two-sample t-test was performed. Multiple test correction was done by Benjamini-Hochberg with an FDR of 0.05 by using Perseus (v1.6.0.2) corresponding Tables S2 and S9. The Pearson correlation was based on “valid values” for each pRCC type in Perseus.

For comprehensive proteome data analyses, gene set enrichment analysis (GSEA, v3.0) (69) was applied in order to see, if *a priori* defined sets of proteins show statistically significant, concordant differences between pRCC and kidney tissues. Only proteins with valid values in at least seven of ten samples in at least one group with replacing missing values from the normal distribution for the other group were used (Dataset S2). GSEA default settings were applied, except that the minimum size exclusion was set to 5 and KEGG v5.2 was used as a gene set database. The cut-off for significantly regulated pathways was set to a p-value ≤ 0.01 and FDR ≤ 0.10.

For protein-protein interaction (PPI) network analyses, the software tool String v.10.5 has been used to visualize networks of significantly up- or down-regulated proteins with a confidence level of 0.7 (70). High blood contamination was identified in the following five samples: pRCC type I kidney 7 and case 5; pRCC type II case 7; pRCC type IIM kidney 3 and case 2; which were then excluded from further proteome and metabolome analysis. These exclusions were based on the individual GSEA pathway and the String PPI network results, where the pathways “coagulation cascade” or “blood particles” were significantly enriched.

## ACKNOWLEDGMENTS

This work is part of the doctoral dissertation of A.A. Our work is supported by the Max Planck Society and the Foundation for Urologic Research to A.A., A.R., and K.J.

## Author’s Contributions

Proteome profiling was performed by A.A. and V.P., metabolome profiling, data analysis and preparation of figures by A.A; J.F.B. and A.R. recruited pRCC cases, E.K. and S.V. validated histological samples, B.T. performed WES, R.C. and M.A. analyzed WES data; D.M. wrote the manuscript, and conceived and directed the project, A.R. and K.J. reviewed the manuscript.

## Ethics Approval and Consent to Participate

The study was approved by the institutional Ethics Committee (no. EA1/134/12, Charité – Universitätsmedizin Berlin) and was carried out in accordance with the Declaration of Helsinki. All participants gave informed consent.

## Availability of Data and Materials

The datasets generated during the current study are available as supplementary files and in the following repositories:

WES files can be accessed via: https://www.ncbi.nlm.nih.gov/sra; SRA accession number: SUB5437563

Proteomics data via PRIDE: https://www.ebi.ac.uk/pride PXD013523

Metabolomics data via PeptideAtlas: http://www.peptideatlas.org/PASS/PASS01368

