## Supplemental Information for "Papillary renal cell carcinomas rewire glutathione metabolism and are deficient in anabolic glucose synthesis"

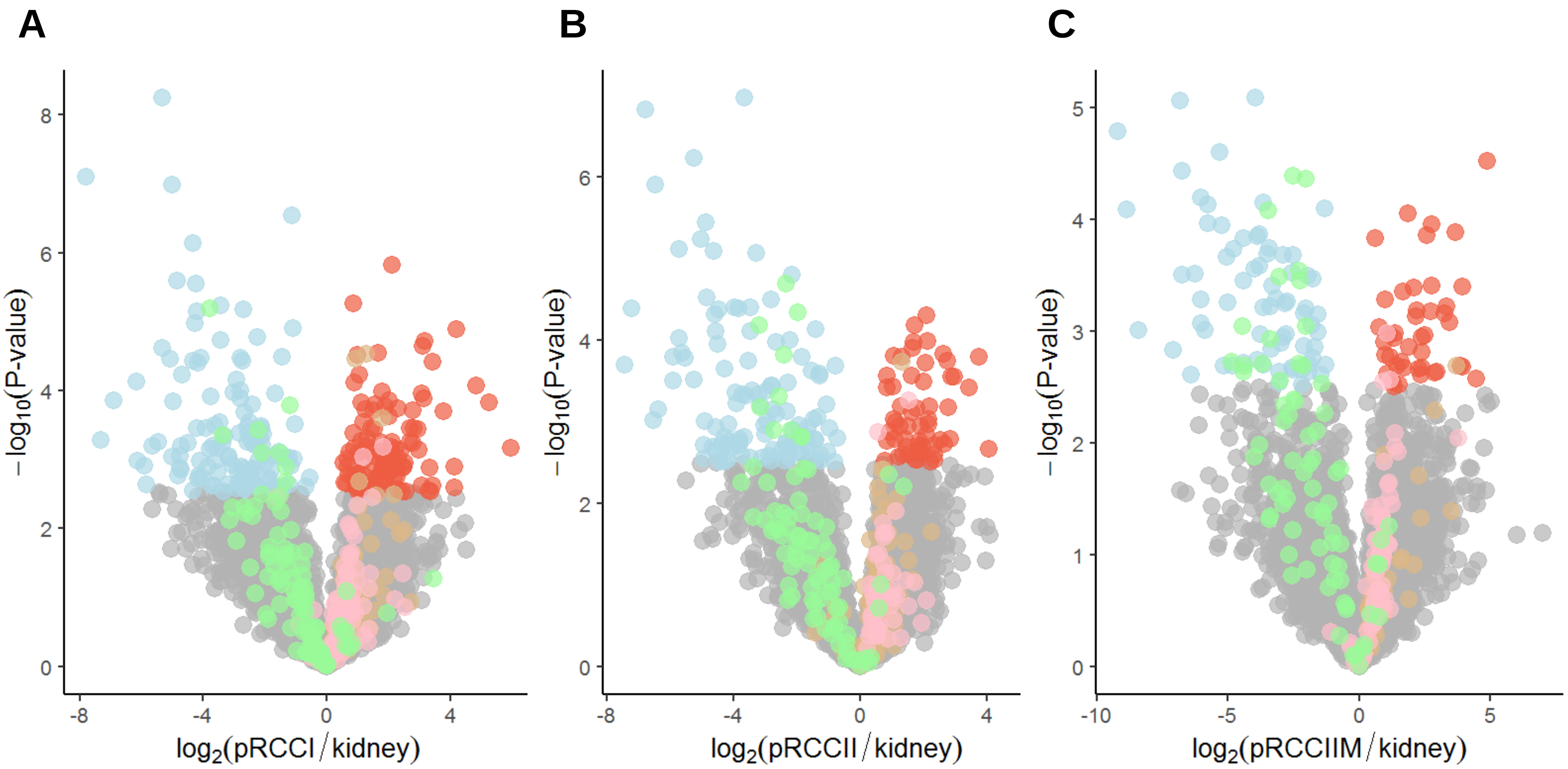
A
B
C
$-\log_{10}$(P-value)
$\log_2$(pRCCI / kidney)
$\log_2$(pRCCII / kidney)
$\log_2$(pRCCIIM / kidney)

**Figure S1. Volcano plot of $log_2$ abundance ratios of (A) pRCC type I, (B) pRCC type II, and (C) pRCC type IIM versus kidney tissues against the $-log_{10}$ (p-value) of the proteome.** Indicated are following pathways: OXPHOS related metabolites in green, and ribosomes in orange. Blue, significantly decreased; red significantly increased in pRCC after considering the Benjamini-Hochberg correction (FDR of 0.05) for multiple testing. The entire list is given in Table S2.

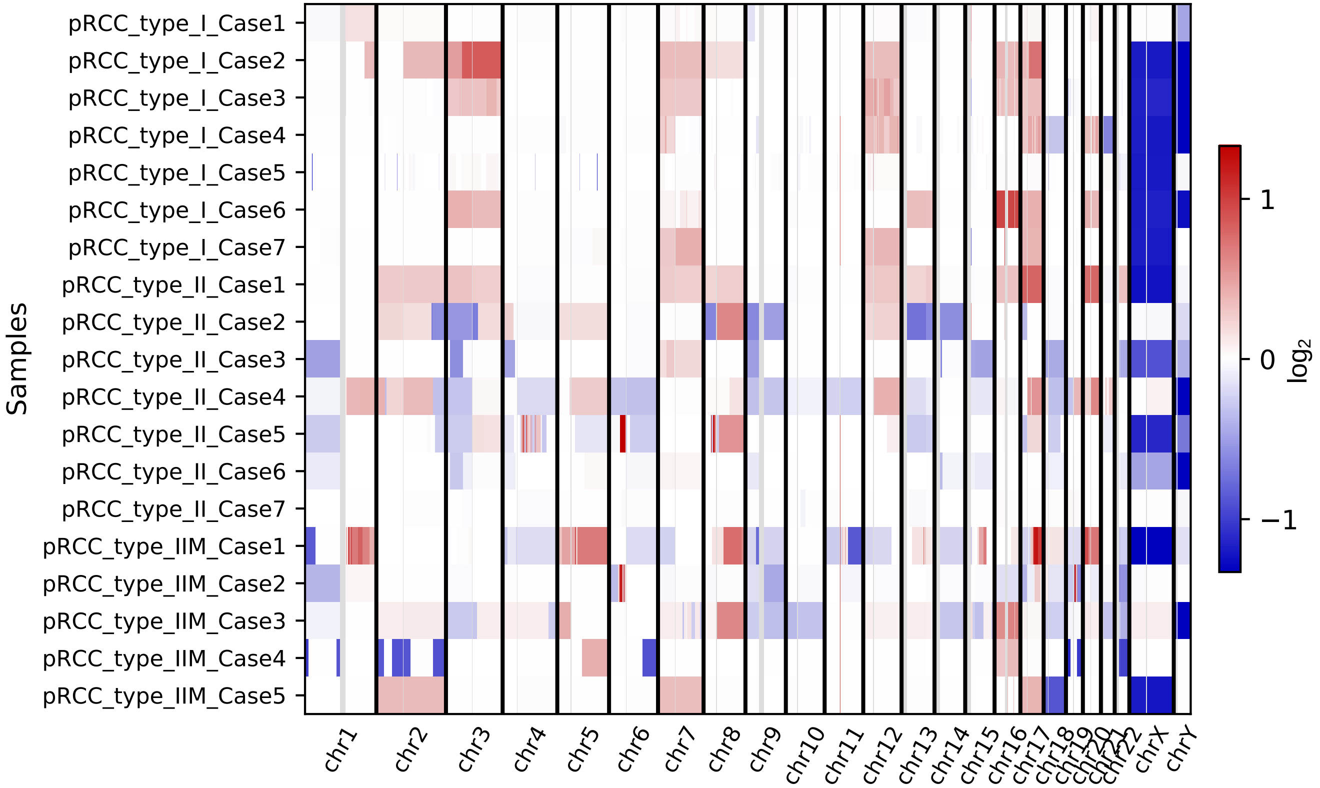

Samples
pRCC_type_I_Case1
pRCC_type_I_Case2
pRCC_type_I_Case3
pRCC_type_I_Case4
pRCC_type_I_Case5
pRCC_type_I_Case6
pRCC_type_I_Case7
pRCC_type_II_Case1
pRCC_type_II_Case2
pRCC_type_II_Case3
pRCC_type_II_Case4
pRCC_type_II_Case5
pRCC_type_II_Case6
pRCC_type_II_Case7
pRCC_type_IIM_Case1
pRCC_type_IIM_Case2
pRCC_type_IIM_Case3
pRCC_type_IIM_Case4
pRCC_type_IIM_Case5
chr1
chr2
chr3
chr4
chr5
chr6
chr7
chr8
chr9
chr10
chr11
chr12
chr13
chr14
chr15
chr16
chr17
chr18
chr19
chr20
chr21
chr22
chrX
chrY
1
0
−1
$log_2$

**Figure S2. Exome-based copy number variation analysis in pRCC.**

**Table S1. MaxQuant output file featuring the proteome profiles of pRCC I, pRCC II, and pRCC IIM with LFQ intensities.**

**Table S2. Significantly regulated proteins between pRCC I, pRCC II, and pRCC IIM and healthy kidneys.** List of protein groups and $Log_2$ LFQ intensities identified in at least three samples of the kidney and the according pRCC type. Significance was tested with a two-sample t-test and additionally with multiple test correction by Benjamini-Hochberg (FDR 0.05), indicated by a + sign.

**Table S3. The percentage of reconstructed genome covered by the assembly, the mean coverage depth, the number of contigs obtained and the best-predicted haplogroup are reported for each sample**. TT = pRCC, NT = normal tissue, same sample numbers are derived from the same patient. The case with no mitochondrial haplogroup match between pRCC and control tissue is highlighted in yellow.

**Table S4. Identified pathogenic somatic and germline mtDNA mutations.** Indicated are variant alleles, homoplasmy, nucleotide variability, amino acid change, amino acid variability, and the disease score.

**Table S5. Pathway enrichment analysis (GSEA) of proteins between pRCC I, pRCC II, pRCC IIM, and adjacent kidney tissues**. Proteins with a minimum of three valid values in at least one group were used, missing values were replaced from normal distribution. KEGG and Reactome databases were used; the significance threshold was set to ≤0.05 for *p-value* and ≤0.10 FDR, respectively. Significant KEGG pathways were highlighted in orange.

**Table S6. Significantly regulated transcripts between pRCC and healthy kidneys.** Data were retrieved from TCGA, missing values in one group were replaced from normal distribution. Significance was tested with a two-sample t-test and additionally with multiple test correction by Benjamini-Hochberg (FDR 0.05), indicated by a + sign. Expression values are shown as $\log_2$ values. The TCGA sample file names encode the specimen entities as following: controls have 11 as the two last digits, pRCC type I has 01 and type II 05.

**Table S7. Pathway enrichment analysis (GSEA) of transcriptome data between pRCC and healthy kidneys.** KEGG and Reactome databases were used; the significance threshold was set to ≤0.05 for the *p-value* and ≤0.10 FDR, respectively. Significant pathways are highlighted in orange.

**Table S8. Significantly regulated metabolites between pRCC I, pRCC II, and pRCC IIM and healthy kidneys as $\log_2$ peak areas.** Significance was tested with a two-sample t-test and additionally with multiple test correction by Benjamini-Hochberg (FDR 0.05), indicated by a + sign. Cell culture tracing experiments to monitor glutathione and proline *de novo* synthesis. Shown are normalized peak areas of the first transition.

**Table S9. Mass spectrometry settings for targeted metabolite profiling. List of all metabolites with masses, MS conditions, and MRM ion ratios for the targeted LC/MS metabolite approach.** Parameters for detection were optimized with prior measurements of the pure metabolites. If the pure substance was not available, parameters were created via database information (METLIN/HMDB), in-house solutions (MS/MS) or based on values of similar chemical structure (MRM's not tuned in). For metabolites without a detectable third transition, a pseudo-MRM, measuring the precursor mass was used. A retention time (RT) of 0 min indicates that the metabolite was measured continuously, due to large peak widths or lack of optimized elution time information. Added internal standards are shown and the standard used for normalization within the method is highlighted as such. (DP = Declustering Potential, CE = Collision Energy, CXP = Collision Exit Potential).
